# The evolution of nesting behaviour in *Peromyscus* mice

**DOI:** 10.1101/177782

**Authors:** Caitlin L. Lewarch, Hopi E. Hoekstra

**Affiliations:** Department of Organismic and Evolutionary Biology, Museum of Comparative Zoology, Howard Hughes Medical Institute, Harvard University, Cambridge, MA 02138 USA; Department of Molecular and Cellular Biology, Museum of Comparative Zoology, Howard Hughes Medical Institute, Harvard University, Cambridge, MA 02138 USA

**Keywords:** behavioural evolution, comparative method, deer mice, extended phenotype

## Abstract

Structures built by animals, such as nests, often can be considered extended phenotypes that facilitate the study of animal behaviour. For rodents, nest building is both an important form of behavioural thermoregulation and a critical component of parental care. Changes in nest structure or the prioritization of nesting behaviour are therefore likely to have consequences for survival and reproduction, and both biotic and abiotic environmental factors are likely to influence the adaptive value of such differences. Here we first develop a novel assay to investigate interspecific variation in the nesting behaviour of deer mice (genus *Peromyscus*). Using this assay, we find that, while there is some variation in the complexity of the nests built by *Peromyscus* mice, differences in the latency to begin nest construction are more striking. Four of the seven taxa examined here build nests within an hour of being given nesting material, but this latency to nest is not related to ultimate differences in nest structure, suggesting that the ability to nest is relatively conserved within the genus, but species differ in their prioritization of nesting behaviour. We also find that latency to nest is not correlated with body size, climate, or the construction of burrows that create microclimates. However, the four taxa with short nesting latencies all have monogamous mating systems, suggesting that differences in nesting latency may be related to social environment. This detailed characterization of nesting behaviour within the genus provides an important foundation for future studies of the genetic and neurobiological mechanisms that contribute to the evolution of behaviour.

## INTRODUCTION

Animal architectures – from the webs spun by spiders to the dams built by beavers – can both facilitate the study of behaviour and provide insight into the selective forces at that act on behavioural variation (Hansell, 1984, 2005). Such structures can be considered “extended phenotypes,” or traits influenced by genetics but *extended* outside the body of the individual organism (Dawkins, 1982). Building behaviours are often innate and species-specific; for example, the resulting structures have been used for classification purposes in insects and some birds (Hansell, 1984; Knerer et al., 2012; Schmidt, 1964; Winkler et al., 1993). These structures reflect stereotyped patterns of behaviour and the neural circuits that generate these motor patterns, allowing us to study behaviour and the nervous system by proxy. Moreover, the structures themselves serve important functions and can confer readily quantifiable fitness benefits on the animals that construct them (Hayward, 1965; Mainwaring et al., 2014; Sealander, 1952).

A widespread and important type of building behaviour is the collection and processing of environmental materials to produce a nest. Nests serve a wide variety of purposes for the animals that construct them. For small-bodied animals, such as rodents, nests provide insulation and reduce the energy expended on the maintenance of body temperature (Pearson, 1960; Sealander, 1952; Vogt & Lynch, 1982). In animals with altricial young, like many birds and rodents, nests are especially critical to protect offspring from heat loss and predation (Bult et al., 1997; Collias, 1964; Lynch & Possidente, 1978; Southwick, 1955). The nest may even serve as a catalyst for social behaviour — nest and bower construction can be integral to courtship in birds (Mainwaring et al., 2014), and investment in elaborate nests likely has been instrumental in the evolution of eusociality in insects (Hansell, 2005). Depending on the species in question and the environment in which they live, nests may be built in trees, in pre-existing cavities, or in burrow systems that are also constructed by the animal (Collias, 1964; Dooley & Dueser, 1990; Weber & Hoekstra, 2009). While the excavation of burrows is itself a type of animal architecture, nests are often separate structures, made by collecting and processing vegetation and other material from an animal’s environment.

Both the structure of a completed nest and the timing of nest building may be relevant traits for natural selection, and each has distinct implications for the proximate and ultimate factors that contribute to behavioural differences among taxa. Variation in nest structure, as is observed in birds, suggests that animals may differ either in their ability to construct nests or in the desired properties of their nests (Mainwaring et al., 2014). At the level of proximate mechanism, variation could result from morphological differences in the animals, fundamental changes in their stereotyped motor patterns, or changes in a more abstract encoding of the animal’s target structure. Moreover, variation in nest structures suggests that the characteristics of the nest itself have fitness consequences. Prime examples of such relationships include the pendulous entrances of some weaverbird nests, which are protective against snake predation (Collias, 1964; Crook, 1963), or the increased size and weight of robin, warbler, and finch nests built at colder northern versus southern latitudes (Crossman et al., 2011). Variation in the timing of nesting behaviour, on the other hand, implies that animals differ in their motivation to engage in otherwise conserved behavioural patterns, and suggests that the prioritization of nesting relative to other elements of the animal’s behavioural repertoire is relevant for selection. Prioritization can occur at different scales, from time invested over the course of a single night to relative time spent on the behaviour during different seasons. As the collection of nesting material can be energetically costly and expose the animal to predation (Collias, 1964; Mainwaring et al., 2014), it may be beneficial for an animal to prioritize other behaviours in environmental conditions where heat loss, for example, is not a pressing concern. While population differences in nest size have been studied within and between species of rodents (King et al., 1964; Lynch, 1992), we do not know how the prioritization of nesting behaviour has evolved.

To determine how and why these features of nesting behaviour evolve, we focused on deer mice (genus *Peromyscus*), which have adapted to a wide range of habitats and microhabitats across North America (Bedford & Hoekstra, 2015; Blair, 1950; Dewey & Dawson, 2001). Specifically, deer mice live in climates with pronounced differences in winter temperatures (King et al., 1964), vary in body size, a trait associated with adaptation to cold in other rodents (Lynch, 1992), and have distinct social behaviour and parental care (Jašarević et al., 2013; Turner et al., 2010), all of which may affect nestbuilding behaviour. Importantly, while these species have evolved in different environments, laboratory colonies allow us to perform behavioural experiments under carefully controlled conditions using animals that share a common environment (Bedford & Hoekstra, 2015). This is therefore an opportunity to explore the evolutionary consequences of different environmental parameters on heritable variation in nest-building behaviour.

Here we develop a novel behavioural assay to evaluate natural variation in both ability and motivation to nest in seven species and subspecies of *Peromyscus* mice. This detailed characterization of thermoregulatory nesting behaviour then provides a foundation to understand the evolution of this behaviour in natural populations.

## METHODS

### Ethical Note

All experimental procedures were approved by the Harvard University Institutional Animal Care and Use Committee. The animal housing facility in which these tests were performed maintains full AAALAC accreditation.

### Experimental Cohort

We selected adult, reproductively inexperienced animals of both sexes from seven laboratory colonies of *Peromyscus*, representing five species, with well-characterized ecology and social systems (Table 1). While these colonies were isolated from natural populations (brought in from the wild between 2 and 71 years ago, depending on strain; Table 1), all animals in this study were born in captivity.

**Table 1:** Experimental Cohort

| Species<br>(common name) | County Isolated | Year in<br>Captivity* | Sample Size<br>**total** (males, females) | Avg. Weight,<br>Males<br>(g ± sd) | Avg. Weight,<br>Females<br>(g ± sd) | Avg. Age<br>(days ± sd) |
| --- | --- | --- | --- | --- | --- | --- |
| *P. maniculatus nubiterrae*<br>(cloudland deer mouse) | Westmoreland<br>County, PA | 2010 | **47** (31,16) | 18.7 ± 2.3 | 15.8 ± 2.8 | 164 ± 184 |
| *P. maniculatus bairdii*<br>(deer mouse) | Washtenaw County,<br>MI | 1946-1948 | **95** (62,33) | 20.3 ± 3.5 | 16.9 ± 1.6 | 106 ± 50 |
| *P. polionotus subgriseus*<br>(oldfield mouse) | Marion County, FL | 1952 | **130** (80,50) | 14.3 ± 1.9 | 15.5 ± 1.7 | 107 ± 57 |
| *P. polionotus leucocephalus*<br>(Santa Rosa Island beach<br>mouse) | Okaloosa County, FL | 2015 | **37** (23,14) | 14.2 ± 1.1 | 14.4 ± 2.7 | 71 ± 17 |
| *P. leucopus*<br>(white-footed mouse) | Avery County, NC | 1982-1985 | **35** (22,13) | 21.7 ± 4.1 | 20.1 ± 2.7 | 66 ± 7 |
| *P. gossypinus*<br>(cotton mouse) | Jackson County, FL | 2009 | **27** (19,8) | 25.2 ± 7.0 | 21.9 ± 3.3 | 72 ± 9 |
| *P. californicus*<br>(California mouse) | Ventura County, CA | 1979-1987 | **48** (25,23) | 42.1 ± 5.2 | 41.5 ± 7.0 | 126 ± 30 |

\*Some species were brought into captivity multiple times over several years, see (Bedford & Hoekstra., 2015). Female *maniculatus bairdii* animals give birth to their first litter when they are approximately three months old (Bedford & Hoekstra, 2015), and generation times for other species are similar in the lab.

### Animal Husbandry

All animals were bred and maintained under the same controlled conditions. We kept the animal housing rooms on a 16:8 LD cycle at 22°C. We housed animals in ventilated polysulfone mouse cages (Allentown, NJ) of standard size (19.7cm wide × 30.5cm long × 16.5cm high), with the exception of the *P. californicus* animals, which were housed in rat cages (28.6cm wide × 39.4cm long × 19.3cm high) due to their large body size (Allentown, NJ). For ordinary housing, we provided all cages with 2.5g of compressed cotton “Nestlet” (Ancare, Bellmore, NY), 8-10g folded paper “Enviro-Dri” nesting material (Shepherd Specialty Papers, Watertown, TN), a 0.6cm layer of Anderson’s Bed-o-cob (The Andersons, Inc., Maumee, OH), and enrichment consisting of a red polycarbonate (9.5cm × 4.8cm × 7.6cm) mouse hut (BioServ, Flemington, NJ) or a 15.2cm × 7.6cm inside diameter rat tunnel for the large *P. californicus* animals (BioServ, Flemington, NJ). All animals had *ad libitum* access to water and irradiated LabDiet Prolab Isopro RMH 3000 5P75 (LabDiet, St. Louis, MO). We socially housed animals in groups of 2-5 by species and sex after weaning (23 days for most species, 30 days for *P. californicus*), then tested them as adult virgins, averaging 2-6 months old (Table 1).

### Behavioural Paradigm

#### Standard Behavioural Assay

Nesting behaviour in rodents is often assessed by measuring the weight of nesting material an animal uses over 24 hours (Hartung & Dewsbury, 1979; King et al., 1964; Layne, 1969; Lynch & Hegmann, 1973), which is readily quantifiable but can obscure variation in the timing of the behaviour or the structure of the nests the animals construct. To measure these aspects of nesting behaviour, we designed a novel assay that consists of an overnight habituation period followed by three consecutive days of testing. On the day before a trial began, we weighed and singly housed adult virgin animals in new mouse cages (including *P. californicus*) with 5g of compressed cotton nesting material (or two “nestlets”, see above), 0.6cm layer of Anderson’s Bed-o-cob, and a red polycarbonate mouse hut. On the morning following habituation to the novel cage, we took photos of the nest from up to three angles (top and two side views), then removed the mouse hut and replaced all cotton nesting material with 5g of fresh compressed cotton nestlet. The replacement of nesting material during these trials always occurred between 4.5 and 6.5 hours after the lights came on. At one hour after the replacement of nesting material, we again took photographs of the nest from multiple angles and added the mouse hut back to the cage. We repeated this process on the following two mornings for a total of three sets of photographs (day 1, day 2, and day 3) at each of the two time points (1h and overnight). Research assistants blinded to the species and sex of the animal later scored these nest photographs according to a standardized scale (Fig. 1; Supplemental Table S1). Scores ranged from 0 (no visible shredding) to 4 (a full “dome” nest with overhead coverage) with only full and half scores given.

**Figure 1:**
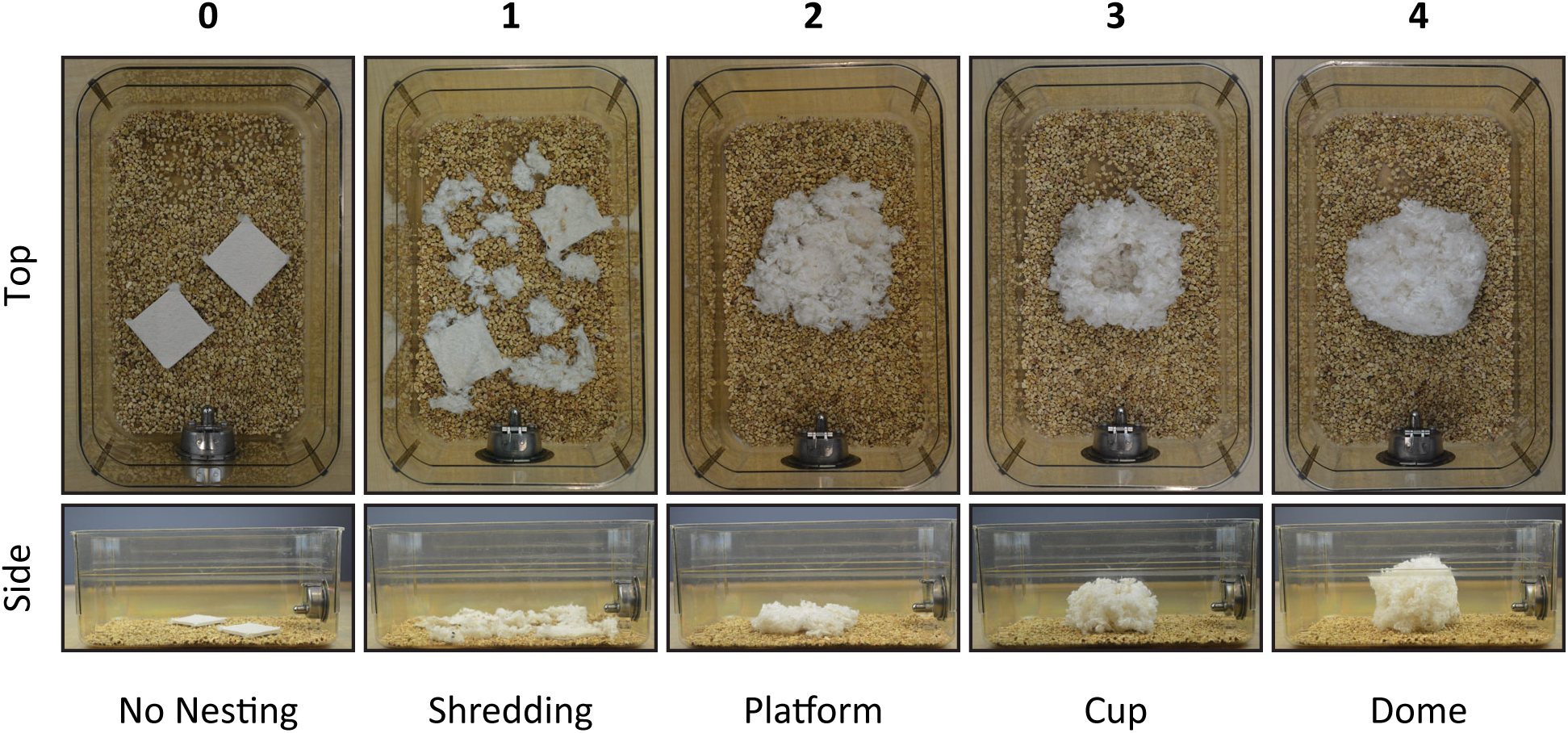
Nest scoring scale. Nests are scored on a scale from 0 (no manipulation of the nesting material) to 4 (a full cotton “dome”) in increments of 0.5. Representative nests for each of the five integer scores are shown from both the top and side view, and a brief descriptor is provided for each. Classification is according to criteria provided in Supplementary Table S1.

#### Increased Nesting Material

To examine whether the amount of nesting material had an impact on nest scores in the largest species (*P. californicus;* approximately 42g, on average), we modified the nesting experiment in two ways. First, we singly housed an independent cohort of 21 adult *P. californicus* animals as above, but provided them with an increasing amount of cotton nesting material on four consecutive days: 5g on day 1, 10g on day 2, 15g on day 3, and 20g on day 4. We photographed nests and exchanged cotton nesting material once every 24 hours, and a research assistant blind to experimental conditions scored these photographs as above to establish whether this increase was sufficient to alter overnight nest scores. Based on the results of these experiments, we then assayed an independent group of 23 adult *P. californicus* animals to evaluate their overnight nesting behaviour using 20g of cotton nesting material in an otherwise standard nesting assay (see “Standard Behavioural Assay” above).

### Climate Data

We drew average winter (December/January/February) temperature data from National Oceanic and Atmospheric Administration (NOAA) 30-year climate normals (Arguez et al., 2010), and averaged these data by state or county of origin for each colony (Table 2).

**Table 2:** Environmental Context

| Taxon | Latitude<br>(°N)<sup>A</sup> | Avg.<br>Winter<br>Temp.<br>(°C)<sup>B</sup> | Habitat | Habitat<br>Use | Habitat Ref. | Burrows<sup>C</sup> | Nest Location | Mating<br>System<sup>D</sup> |
| --- | --- | --- | --- | --- | --- | --- | --- | --- |
| *P. m. nubiterrae* | 40.2 | -1.2 | forest | semi-arboreal | (Blair, 1950) | simple/short | arboreal; tree cavities (Wolff & Durr, 1986; Wolff & Hurlbutt, 1982) | M |
| *P. m. bairdii* | 42.3 | -3.4 | prairie/grassland | terrestrial | (Blair, 1950) | simple/short | in burrows (Morris & Kendeigh, 1981) | P |
| *P. p. subgriseus* | 29.2 | 14.4 | sandy soil/grassland | semi-fossorial | (Blair, 1950) | complex/long | in burrows (Dawson et al., 1988) | M |
| *P. p. leucocephalus* | 30.4 | 11.0 | white sand beach | semi-fossorial | (Blair, 1950; Sumner, 1926) | complex/long | in burrows (Blair, 1951) | M |
| *P. leucopus* | 36.1 | 0.9 | deciduous forest | semi-arboreal | (Blair, 1950; Lackey et al., 1985) | simple/short | seasonally dependent; arboreal or ground nests (Wolff & Durr, 1986; Wolff & Hurlbutt., 1982) | P |
| *P. gossypinus* | 30.7 | 11.3 | hardwood forest, mesic hammocks, swamps | semi-arboreal | (Wolfe & Linzey, 1977) | no laboratory data | diverse but arboreal preferred; in or under logs and stumps, in tree cavities (Ivey, 1949; Klein & Layne, 1978; Wolfe & Linzey, 1977) | P |
| *P. californicus* | 34.1 | 12.3 | scrub/chaparral | semi-arboreal | (Clark, 1936; M'Closkey, 1976; Merritt, 1978; Meserve, 1977) | none | under logs, in woodrat (*Neotoma*) dens (Merritt, 1974, 1978) | M |

A. Latitude of county of origin (Table 1).
B. 30-year NOAA climate normals for average Dec/Jan/Feb temperatures, pooled by county of origin for each colony (Arguez et al., 2010).
C. Burrows produced in a laboratory assay from (Weber & Hoekstra, 2009), with the exception of *P. m. nubiterrae* (Hu & Hoekstra, 2017).
D. Most likely mating system (Monogamous or Promiscuous) according to (Turner et al., 2010), with the exception of *P. m. nubiterrae* (Wolff & Cicirello, 1991) and *P. gossypinus* (Dewsbury et al., 1980; McCarley, 1959; Pearson, 1953).

### Data Analysis

We performed statistical analyses in R using non-parametric methods for the ordinal nest scores. We summarized an animal’s behaviour across the three trial days by its median score (to reflect central tendency) or its maximum score (to represent best effort) at each time point after the replacement of nesting material. To identify differences between groups, we used Kruskal-Wallis rank-sum tests, and then used Bonferroni-corrected Wilcoxon rank-sum tests for subsequent pairwise comparisons between species or sexes. For the experiment that tests the effects of increased access to nesting material in a cohort of *P. californicus* animals, we used a Friedman rank-sum test.

For comparative analyses, we first generated an ultrametric tree using Grafen’s method (Grafen, 1989; Symonds & Blomberg, 2014) and the known topology of the species relationships (Fig. 2A; Bedford & Hoekstra, 2015; Bradley et al., 2007; Weber & Hoekstra, 2009). To test for a relationship between species-level median 1h nest scores and species-level average weights, latitudes of origin, and winter temperatures at sites of origin, we performed phylogenetic generalized least squares (PGLS) analysis using the ape and nlme packages in R (Paradis et al., 2004; Pinheiro et al., 2017; Symonds & Blomberg, 2014). Covariance due to relatedness was modelled by Brownian motion using the corBrownian function in ape. The covariance was then included as a correlation parameter in the generalized least squares analyses in nlme. The effect of each environmental variable on 1h nest scores was tested independently. To test whether short nesting latency is dependent on other discrete traits (complex burrowing or mating system, as indicated in Table 2), we performed Pagel’s binary character correlation test using the fitPagel function in the phytools package in R (Pagel, 1994; Revell, 2012). For this test, we utilized the fitMk method, allowed all rates of change to be different between states (model=“ARD”), and set nesting latency (short vs. intermediate/long) to be dependent on the state of either mating system (monogamous vs. promiscuous) or burrow complexity (complex vs. simple/absent). As there are no laboratory data on burrowing behaviour in *P. gossypinus*, this species was excluded from the latter analysis.

**Figure 2:**
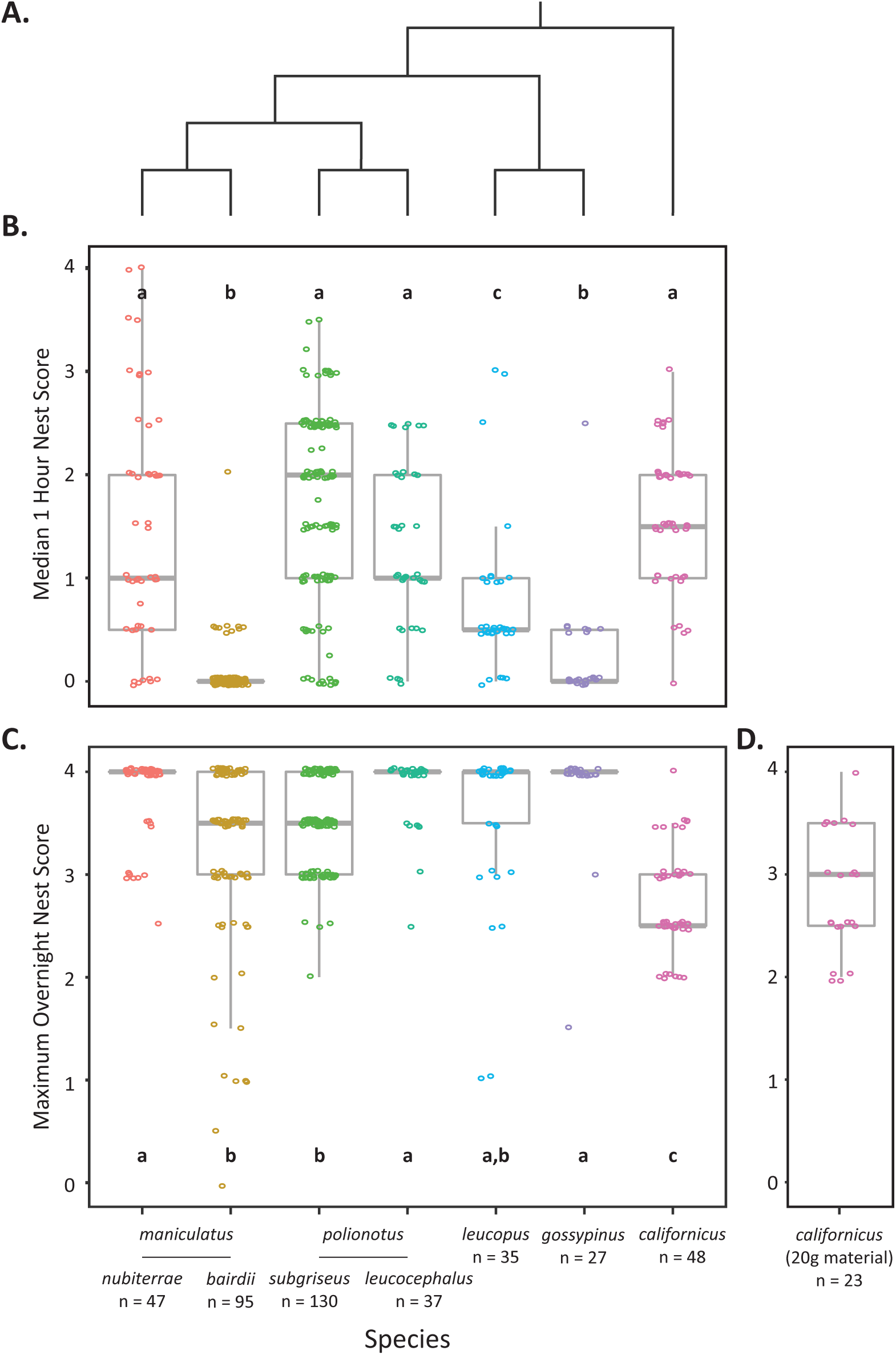
Interspecific differences in nesting behaviour. (A) Phylogenetic relationships among the *Peromyscus* taxa included in this study; modified from (Bedford & Hoekstra, 2015; Weber et al., 2009). (B) Median nest scores 1h after receiving new nesting material and (C) maximum overnight nest scores for each animal over the 3 trial days. Letters indicate species groups that do not significantly differ from one another, while all other pairwise comparisons are significant (Wilcoxon rank-sum test, Bonferroni-corrected *P*<0.05). (D) *P. californicus* animals given 20g of nesting material do not differ in maximum overnight scores from those given 5g of nesting material (Wilcoxon rank-sum test, *P*=0.47). Sample sizes are provided below.

## RESULTS

### Interspecific variation in nesting latency

To measure an animal’s motivation to nest, we assayed individuals from seven *Peromyscus* taxa with known evolutionary relationships (Fig. 2A). First, we analysed the median of the three scores an animal received one hour after the replacement of nesting material, which reflects the tendency of the animal to begin nesting shortly after their nest is disturbed. Scores at 1h were significantly correlated across the three days in the full dataset (Spearman rank correlations: day 1 vs. day 2 *r*_*s*_ = 0.75, day 1 vs. day 3 *r*_*s*_ =0.68, day 2 vs. day 3 *r*_*s*_ = 0.78, *N*=419, *P*<2.2×10^-16^ for each), and species comparisons were largely the same whether three-day medians or maxima were used (see below). The median nest scores at 1h following the initiation of the trial varied dramatically among the taxa we assayed (Fig. 2B; Kruskal-Wallis test: *H*_6_= 216.85, *P*<2.2×10^-16^). Four taxa (*P. m. nubiterrae*, *P. p. subgriseus*, *P. p. leucocephalus*, and *P. californicus*) had high, statistically indistinguishable scores at the 1h time point (Supplemental Table S2, Wilcoxon rank-sum test, Bonferroni-corrected P>0.05 for each pairwise comparison), suggesting that they began to construct their nests relatively quickly and progressed past the point of just shredding the material. In addition, these four taxa differed significantly from the other three taxa we assayed (Supplemental Table S2, Wilcoxon rank-sum test, Bonferroni-corrected p<0.05 for each pairwise comparison). *P. leucopus* animals received intermediate scores that reflect a tendency to shred the material, but not arrange it into a nest, at the 1hr time point. These scores were significantly different from those received by all other species (Supplemental Table S2, Wilcoxon rank-sum test, Bonferroni-corrected *P*<0.05 for each pairwise comparison). Finally, *P. m. bairdii* and *P. gossypinus* animals had equivalently low scores (Supplemental Table S2, Wilcoxon rank-sum test, Bonferroni-corrected *P*=0.29), which indicate that they did not manipulate the nesting material in the first hour and suggest that they are relatively slow to initiate nest construction. Ranking of each taxon’s performance largely followed the same pattern whether maximum or median nest scores were used (Supplemental Figure S1). The only exception was one species difference: while the median nest scores of *P. gossypinus* and *P. m. bairdii* animals at 1h were indistinguishable, *P. gossypinus* were slightly more likely to shred the nesting material on at least one of the trial days and therefore had slightly, but significantly, higher maximum scores (Wilcoxon rank-sum test: *W*=839, *N*_1_=27, *N*_2_=95, Bonferroni-corrected *P* = 0.049). We note that taxa differ in the variance of their nest scores: this likely results from within-species variation in nest-building efficiency, time spent nesting, and/or the precise initiation time during the first hour. In sum, based on our analysis of 1h median scores, we identified three main groups of nest builders in our assay: those with short, intermediate or long latencies to nest.

### Interspecific differences in nesting ability

We next asked whether these taxa differed in their overall ability to construct a three-dimensional nest. To establish the highest-scoring nest that an animal was capable of producing, we used the maximum score achieved over the individual’s three overnight time points, which represents the animal’s best effort during the longest interval of the trial. Maximum overnight scores varied significantly among taxa (Fig. 2C; Kruskal-Wallis test: *H*_6_=127.21, *P*<2.2×10^-16^), although most animals built full or partial domes. The highest scoring nests were consistently constructed by *P. m. nubiterrae*, *P. p. leucocephalus*, *P. leucopus*, and *P. gossypinus* animals, which tended to build statistically indistinguishable full domes (Supplemental Table S2, Wilcoxon rank-sum test, Bonferroni-corrected P>0.05 for each pairwise comparison). Three taxa – *P. m. bairdii*, *P. p. subgriseus*, and *P. leucopus* – had equivalently high maximum scores (Supplemental Table S2, Wilcoxon rank-sum test, Bonferroni-corrected P>0.05 for each pairwise comparison), and *P. m. bairdii* and *P. p. subgriseus*, which tended to build domes with only partial cover, were significantly different from all but *P. leucopus* animals (Supplemental Table S2, Wilcoxon rank-sum test, Bonferroni-corrected *P*<0.05 for each pairwise comparison). Finally, *P. californicus* animals tended to build nests with walls but without overhead cover, and had significantly lower maximum nest scores than all other species tested (Supplemental Table S2, Wilcoxon rank-sum test, Bonferroni-corrected *P*<0.05 for each pairwise comparison). Notably, we found that all species had at least one individual who constructed a domed nest with full cover (maximum nest score, “4”) during the assay, suggesting that all species are capable of building a “complete” nest if given enough time. However, some species showed a large variance in nest scores, and *P. californicus* tended to have lower maximum scores than the other species.

### Nest-building behaviour in the large *P. californicus* mice

*P. californicus* animals are much larger than the other taxa included in this study (Table 1), and therefore might require more material to construct a dome nest with overhead cover. To test the possibility that these animals built lower-scoring nests because 5g of nestlet was an insufficient amount of nesting material, we conducted two additional experiments. First, we gave a group of *P. californicus* animals increasing amounts of nesting material on four consecutive days and evaluated the nests they produced in each 24-hour interval. We found that increasing nesting material from 5g to 20g could increase overnight nesting scores (Supplemental Fig. S2; Friedman test: *X*^2^_3_=13.468, p=0.004). However, when we provided an independent group of *P. californicus* animals with 20g of nesting material during a three-day trial (Fig. 2D), there was no difference in overnight maximum scores between those *P. californicus* given 5g of nestlet and those given 20g (Wilcoxon rank-sum test: *W*= 609, *N*_1_=23, *N*_2_=48, *P*=0.47). Moreover, the maximum overnight nest scores for *P. californicus* given 20g of nestlet remained significantly lower than the maximum nest scores for all other species (Supplemental Table S2, Wilcoxon rank-sum test, Bonferroni-corrected *P*<0.05 for each pairwise comparison). Thus, the poor nest construction of *P. californicus* in this assay cannot be attributed simply to insufficient nesting material relative to its large body size.

### Sex differences in nesting

We next investigated whether there were any sex differences in the nest scores produced by each species and subspecies. Only two taxa showed evidence of sexual dimorphism in nesting (Fig. 3). Both male *P. m. nubiterrae* and male *P. p. subgriseus* animals built higher scoring nests than their female counterparts one hour after the start of the assay (Supplemental Table S3, Wilcoxon rank-sum test, Bonferroni-corrected *P* = 0.03 and 0.002, respectively), and *P. p. subgriseus* males also built higher scoring nests at the overnight time point (Supplemental Table S3, Supplemental Fig. S3; Wilcoxon rank-sum test, Bonferroni-corrected *P* = 0.008). No other species showed evidence of sex differences in nest scores at either time point (Supplemental Table S3, Wilcoxon rank-sum test, Bonferroni-corrected *P* >0.05 for each pairwise comparison). Therefore, while there was no sexual dimorphism in nesting behaviour for most taxa, in both instances when sex differences were observed, males constructed higher-scoring nests than the females.

**Figure 3:**
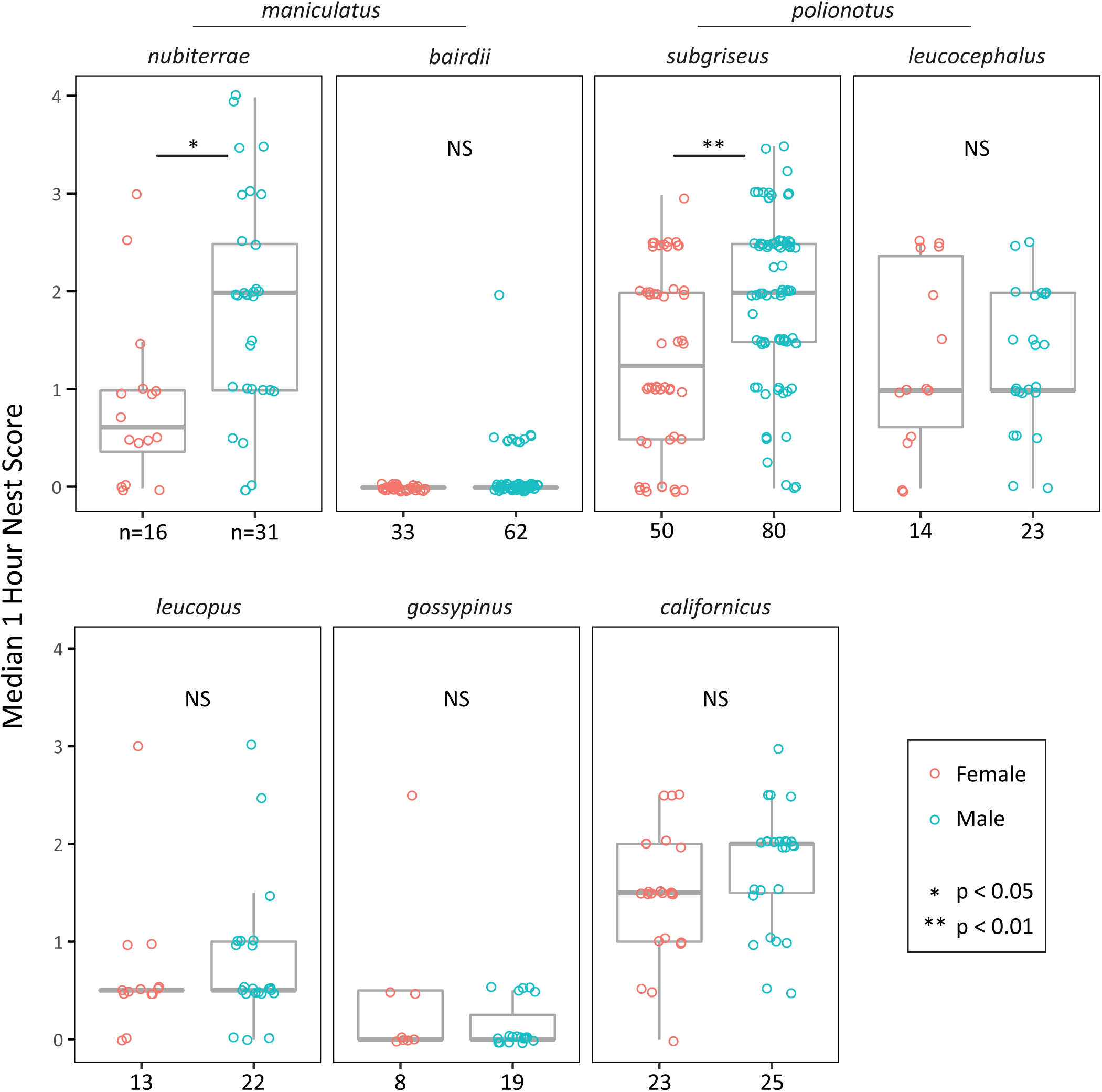
Sex differences in nesting latency. Sex-specific median 1h nest scores for each *Peromyscus* taxa tested. Significant sex differences in median nest score occurred in only two groups: *P. maniculatus nubiterrae* (Wilcoxon rank-sum test, Bonferroni-corrected *P*=0.03) and *P. polionotus subgriseus* (*P*=0.002). Sample sizes are provided below. NS = non-significant.

### Association between body size and nest building

To determine whether body size had an effect on nest-building behaviour, we tested for correlations between weight and performance in the nesting assay. We found that weight significantly varied by species, sex, and species-by-sex interactions in our experimental cohort (two-way ANOVA, main effect of species: *F*_6,352_ =365.9, *P*<2×10^-16^; main effect of sex: *F*_1,352_ = 5.3, *P*=0.02; interaction: *F*_6,352_ =4.1, *P*=0.0005). However, there was no evidence that species-level average weights alter median 1hr nest scores (Fig. 4A, phylogenetic generalized least squares, median 1hr nesting score by average weight: coefficient= -0.01, SE=0.06, *t*=-0.15, *P*=0.89). Likewise, when we divided the animals by species and sex, we found no correlation between weight and median nest score at 1h (Supplemental Table S4, Spearman’s rank correlations, Bonferroni-corrected *P* >0.05) or maximum overnight nest score (Supplemental Table S4, Spearman’s rank correlations, Bonferroni-corrected *P* >0.05) within any of the species-sex groups. Thus, while average weight varied almost three-fold among species, weight was not associated with nesting behaviour in our assay.

**Figure 4:**
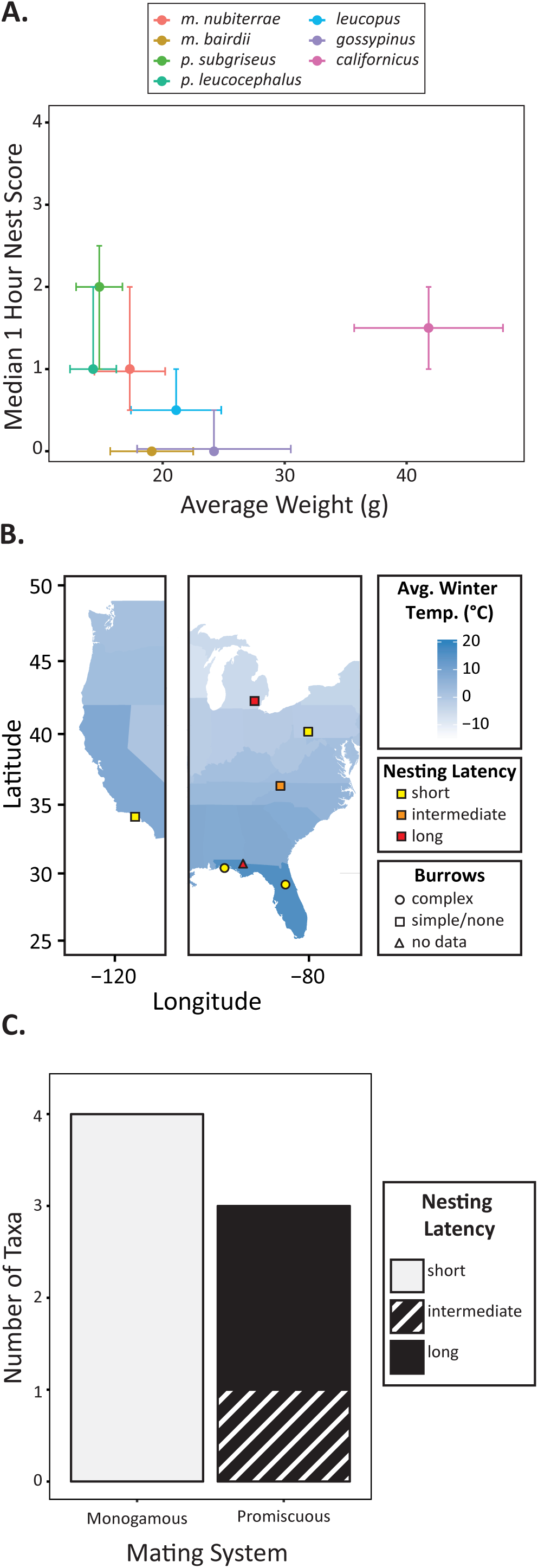
Environmental factors and nesting behaviour. (**A**) Species median 1h nest scores and average weight are plotted with bars indicating interquartile range (nest score) and standard deviation (weight). (**B**) Sites of colony origin (on US map), burrow shape (by symbol), and nesting latency (by colour) are indicated for each taxon (following legend). Map colours represent average winter temperatures by state (Arguez et al., 2010). Nesting latency category (short, intermediate, long) was determined by the significant species groups depicted in Fig. 2B. (**C**) Association between a taxon’s mating system and nesting latency.

### Association between environment and nest construction

We next asked whether there was an association between performance in the nesting assay and several additional environmental covariates, including latitude and average winter temperature of origin, burrow construction, and mating system. Neither latitude nor average winter temperatures were significantly associated with median 1hr scores in these species (Table 2, Fig. 4B; phylogenetic generalized least squares, median 1hr nest score by latitude: coefficient =-0.05, SE= 0.08, *t*=-0.71, *P*=0.51; median 1hr nest score by average winter temperature: coefficient = 0.03, SE= 0.05, *t*=0.69, *p*=0.52). Moreover, nesting latency does not appear to be influenced by burrowing behaviour: a model in which short nesting latency is dependent on building complex burrows does not fit the data significantly better than a model where the two traits are independent (Table 2, Fig. 4B; Pagel’s binary character correlation test: AIC (independent model) = 21.97, AIC (dependent model) = 23.88, likelihood ratio=2.09, *P*=0.35). However, a model in which short nesting latency depends on mating system fits the observed data significantly better than a model assuming the two traits are independent (Table 2, Fig. 4C; Pagel’s binary character correlation test: AIC (independent model)= 26.42, AIC (dependent model)=21.21, likelihood ratio = 9.21, *P*=0.01). With the caveat that the sample size for comparisons among taxa is small, these data suggest that mating system is correlated with nesting latency but the other abiotic environmental factors we examined are not.

## DISCUSSION

Nesting is important for survival in rodents, but it is not clear how this behaviour varies among species or which evolutionary pressures drive these changes. Here we designed a novel high-throughput phenotyping paradigm to evaluate variation in both nest structure and the timing of nesting behaviour in closely related species of deer mice. We found that *Peromyscus* mice are generally able to construct dome-shaped nests, but vary strikingly in their latency to do so. Because nesting latency is not simply correlated with phylogeny, this raises the possibility that natural selection may contribute to inter-taxon variation. When we tested for correlations between latency to nest and several abiotic and biotic variables, we found that mating system, but surprisingly not climate or body size, is correlated with nesting behaviour.

Nesting has been well studied in laboratory models (e.g. Lisk et al., 1969; Lynch, 1980). However, the majority of these nesting experiments, including some studies in *Peromyscus*, measure the amount of nesting material that an animal pulls into its cage or the final nest structure achieved over a 24-hour period (Hartung et al., 1979; King et al., 1964; Layne, 1969; Lynch & Hegmann, 1973). By contrast, we focus on both the timing of the behaviour and the final nest structure. By evaluating nests just one hour after the replacement of nesting material, we are able to assess whether the animals differ in their latency to begin nest construction—what might be considered a baseline motivation to nest. This is complemented by a second measurement at the more permissive overnight time point, which allows us to evaluate whether animals vary in their overall ability to build three-dimensional structures. This novel phenotyping paradigm therefore allows us to distinguish between animals that differ in their motivation to construct nests of similar shape from those that differ in their ability to construct nests.

Using this approach, we find that even closely-related *Peromyscus* species vary dramatically in their latency to begin nesting, while variation in final nest structure is much more modest. This is in contrast to studies of nesting in birds and insects, where the structures of complete, species-typical nests are highly variable (Collias, 1964; Healy et al., 2008; Knerer et al., 2012; Price & Griffith, 2017; Schmidt, 1964), or even burrow construction in *Peromyscus*, where species excavate cavities that significantly differ in size and shape (Hu & Hoekstra, 2017; Weber & Hoekstra, 2009). The relative conservation of nest structure implies that the ability to produce dome-shaped nests is important for most animals in the genus. However, variation in latency to begin nesting suggests that prioritization of the behaviour varies among taxa. These patterns also imply that variation in nesting in these mice is likely due to altered motivation rather than changes in stereotyped motor patterns, morphology, or target nest structure.

All animals were acclimated to and tested in a common environment, specifically at 22°C, which is below the preferred temperatures (Ogilvie & Stinson, 1966) and thermoneutral zones (Layne & Dolan, 1975; Glaser & Lustick, 1975; Hayward 1965) of many *Peromyscus* species. While these taxa may differ in their behavioural response to this thermal environment due to differences in basal metabolic rate or thermoneutral zone, we found no evidence of a correlation between nesting scores at either time point and weight, a trait strongly related to both metabolic parameters in *Peromyscus* (Hayward, 1965; Hill, 1983). It is worth noting that the positive relationship between body weight and nest weight observed in previous studies of rodent nesting (King et al., 1964; Lynch, 1992; Wolfe, 1970) might be at least partially explained by larger animals requiring more material to build equivalently shaped structures. By focusing on the structure of the nest rather than the weight of nesting material used to construct it, we minimize this confounding factor.

Body size aside, it is reasonable to hypothesize that climate could alter this thermoregulatory behaviour. Other studies have suggested that climate (King et al., 1964; Lynch, 1992) and microclimate (Wolfe, 1970) alter the amount of nesting material used by rodents in natural populations. However, we find only modest variation in final nest shape and no evidence for a relationship between nesting latency and average winter temperatures or latitude of origin, which is frequently used as a proxy for temperature. Nor do we find evidence for an association between nesting latency and the construction of elaborate burrows, which function as microclimates and buffer the animals from changes in ambient temperature (Hayward, 1965; Sealander, 1952; Weber & Hoekstra, 2009). Although these colonies have experienced reduced selective pressure while bred in laboratory settings, there is still no evidence of an association between climate/microclimate and latency when we consider only the colonies founded within the past 10 years (Table 1; *gossypinus*, *polionotus leucocephalus*, and *maniculatus nubiterrae*). Given that most of the variation we observe takes the form of prioritization differences rather than changes in nest size or shape, it may be especially necessary to consider the broader behavioural repertoire of these taxa, including biotic factors that could contribute to the motivation to nest.

While we do not observe a simple relationship between nesting behaviour and any of the abiotic factors we examined, we do find an intriguing correlation between social environment and nesting latency. With the caveats that our sample size is relatively low and that the classification of species as monogamous or promiscuous relies on incomplete evidence, our results suggest that mating system and nesting latency are not independent, with all putatively monogamous species having short latencies to nest. It is possible that this reflects a tendency to invest in a home territory that is more beneficial for monogamous animals than for promiscuous ones (Gaulin & FitzGerald, 1988), or that selection for increased paternal care, a hallmark of monogamous mating systems (Kleiman, 1977), might result in increased motivation to nest even in virgin animals. This potential relationship between social behaviour and the prioritization of nesting behaviour underscores the importance of considering both biotic and abiotic environment when investigating the causes of behavioural evolution.

## CONCLUSION

Measurement of extended phenotypes such as nests allows us to study how behaviours evolve within and between species. Here we showed that the ability to nest is relatively conserved in the genus *Peromyscus*, but latency to begin nest construction is highly variable, even between sister species. This suggests that evolution of nesting behaviour in these animals is characterized by differences in the prioritization of an otherwise conserved behavioural pattern. Intriguingly, while abiotic environment cannot explain these species differences in nesting behaviour, we find a correlation between latency to nest and mating system, with monogamous species prioritizing nesting. Finally, as the innate differences in nesting behaviour in *Peromyscus* appear to be largely changes in the motivation to nest, future studies in this system may elucidate genetic and neurobiological mechanisms that lead to differences in motivation to engage in particular behaviours, a topic with implications far beyond nesting behaviour.

## ACKNOWLEDGEMENTS

We would like to thank Harvard undergraduates A. Bielawski, M. Charifson, E. D’Agostino, M. Noriega, and J. Rhodes for blinded scoring of behavioural assays. N. Bedford and C. Hu assisted with the figures. L. Revell and O. Lapiedra provided advice on comparative methods. The Hoekstra lab provided helpful feedback on the experiments and figures, and N. Bedford, E. Hager, D. Haig, C. Hu, B. König, M. Noriega, K. Pritchett-Corning, and K. Turner commented on the manuscript. We would like to thank the Harvard OAR staff for their assistance with animal husbandry.

---

**Supplementary Figure S1:**
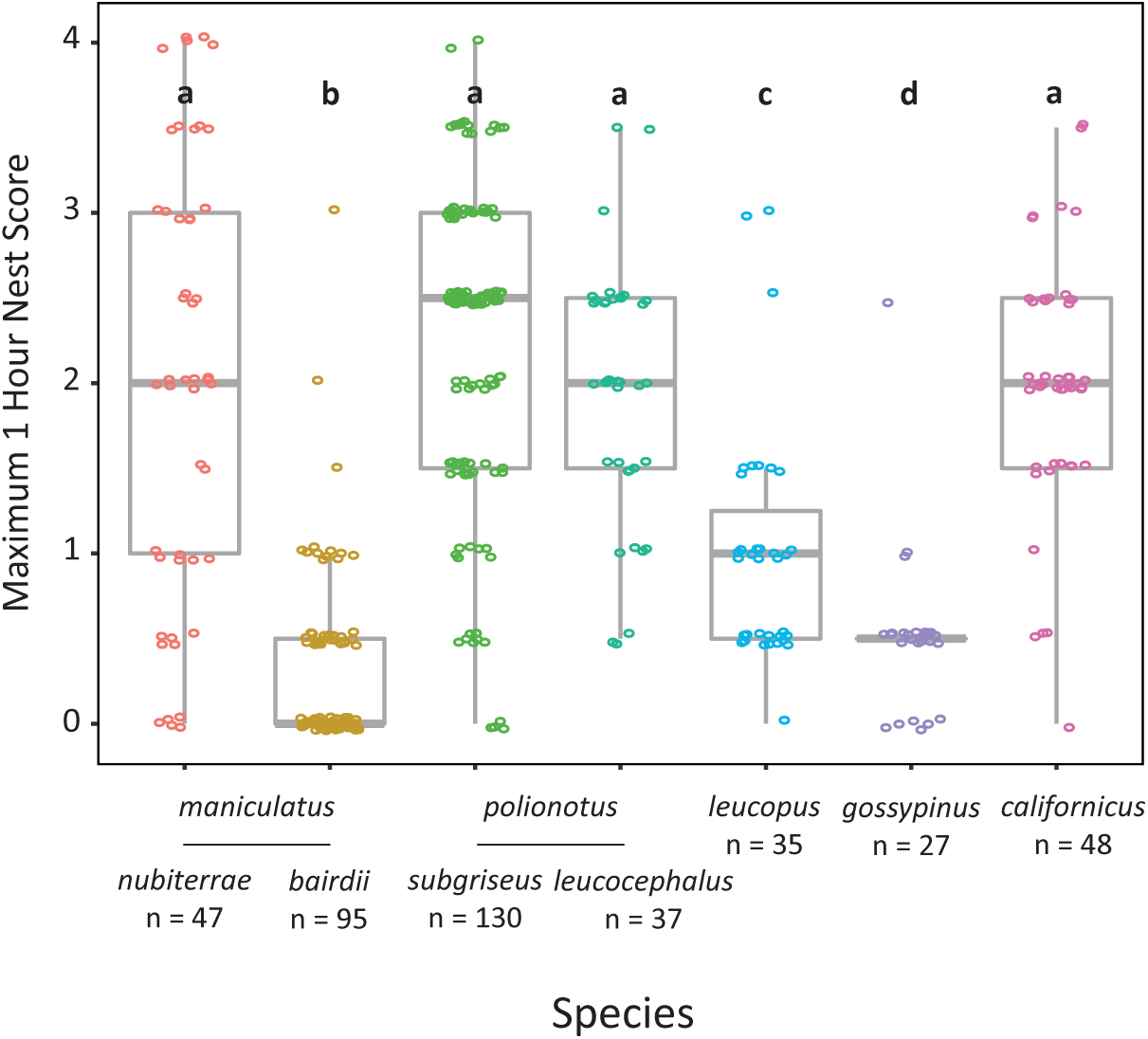
Maximum 1 hour nest scores. Maximum nest score achieved 1h after receiving new nesting material over the 3 trial days. Letters indicate species groups that did not significantly differ from one another, while all other pairwise comparisons were significant (Wilcoxon rank-sum test, Bonferroni-corrected *P*<0.05), largely consistent with median scores (Fig. 2B, Supplemental Table S2), with the exception of a significant difference in maximum scores between *P. gossypinus* and *P. m. bairdii* animals (Wilcoxon rank-sum test, Bonferroni-corrected *P*=0.049). Sample sizes provided below.

**Supplementary Figure S2:**
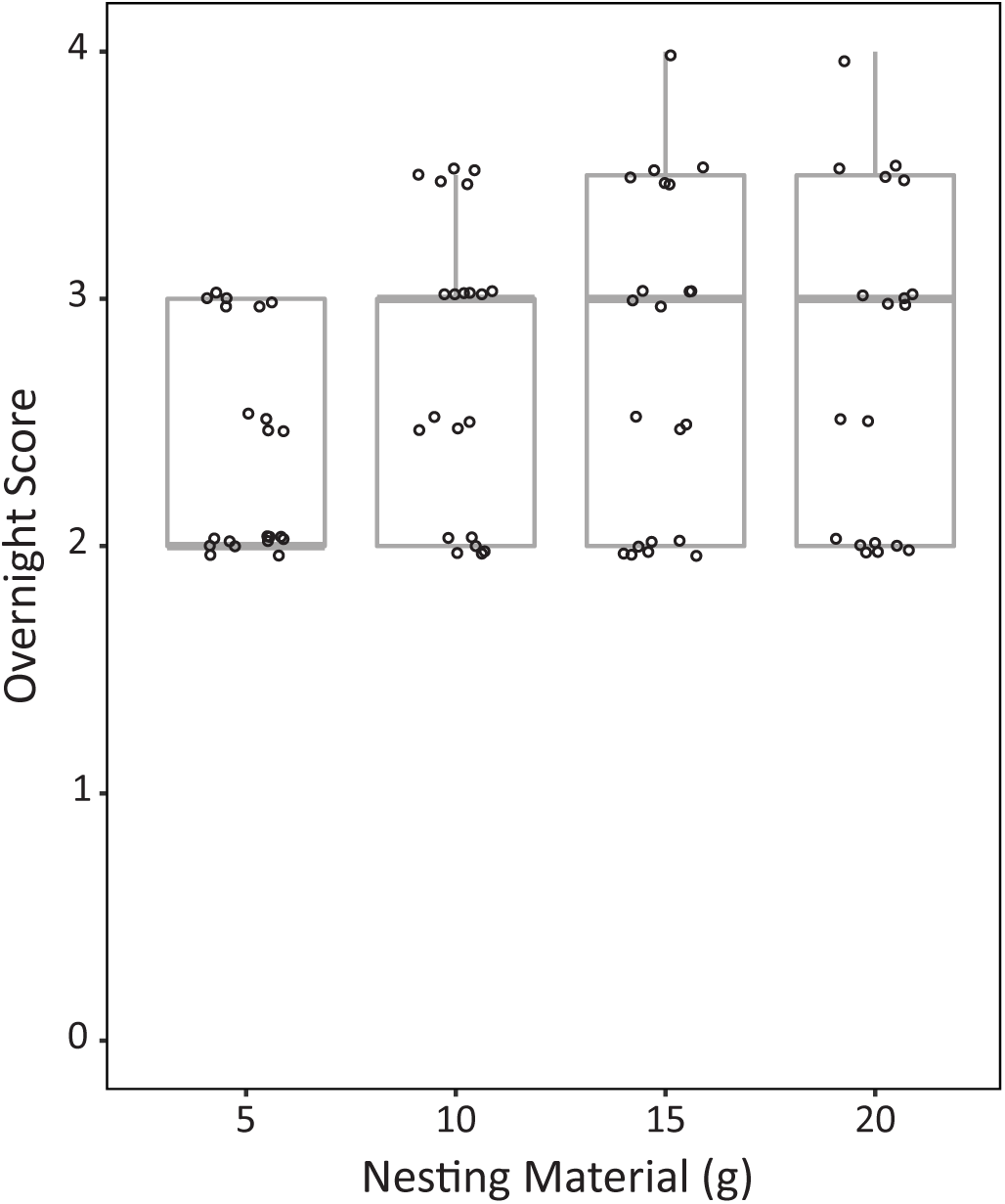
Effect of increasing nesting material in *P. californicus*. Adult animals (*N*=21) were given increasing amounts of nesting material (5g, 10g, 15g and 20g) on 4 sequential days. Higher amounts of nesting material increased overnight nest scores (Friedman repeated measures test, *P*=0.003).

**Supplementary Figure S3:**
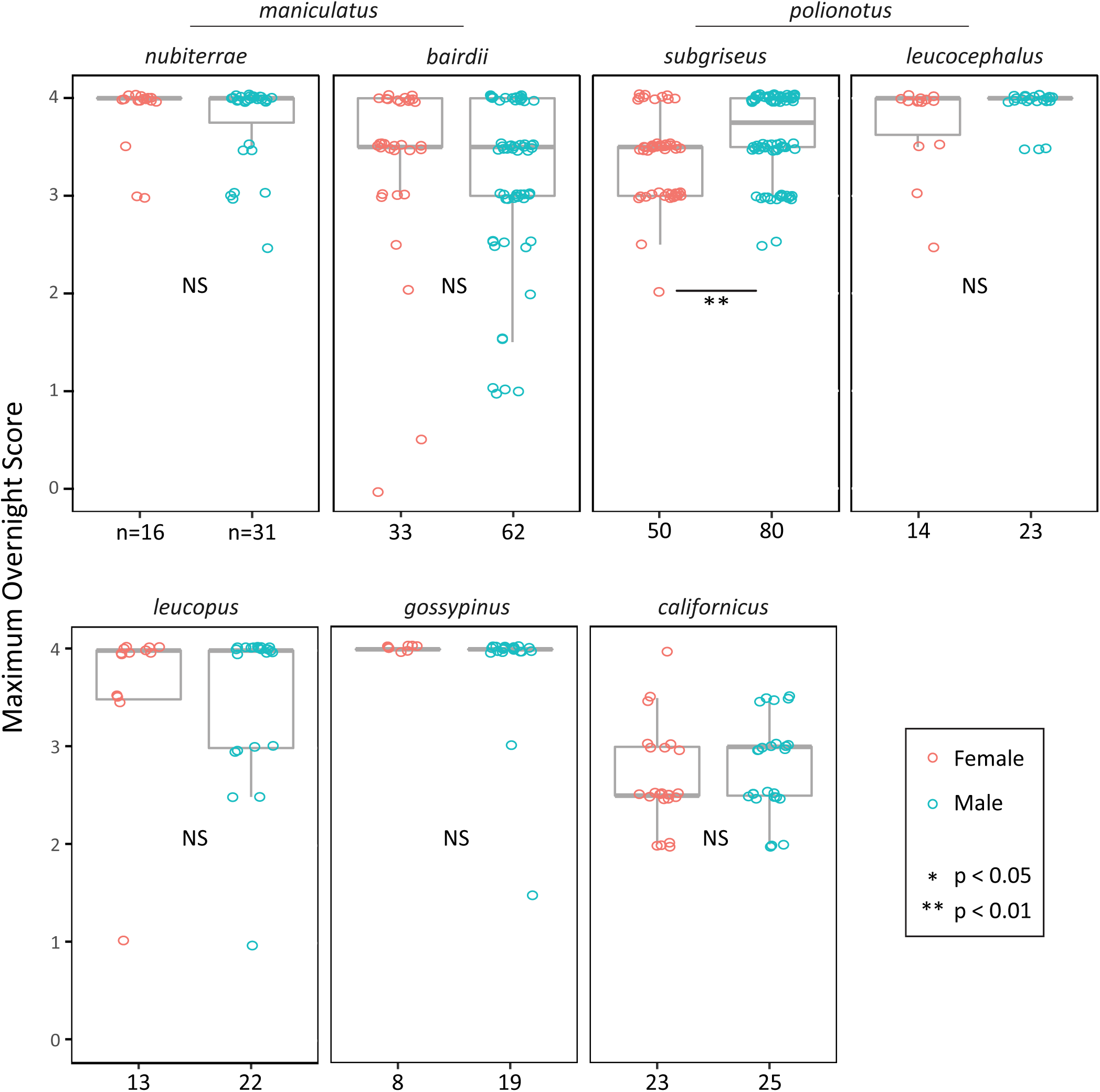
Sex differences in maximum overnight nest scores. Evidence for a significant sex difference overnight nest score occurred in only one taxon: in *P. polionotus subgriseus*, males built higher scoring nests at the overnight time point than females (Wilcoxon rank-sum test, Bonferroni-corrected *P*=0.008). Sample sizes are provided below. NS = not significant.

**Supplementary Figure S4:**
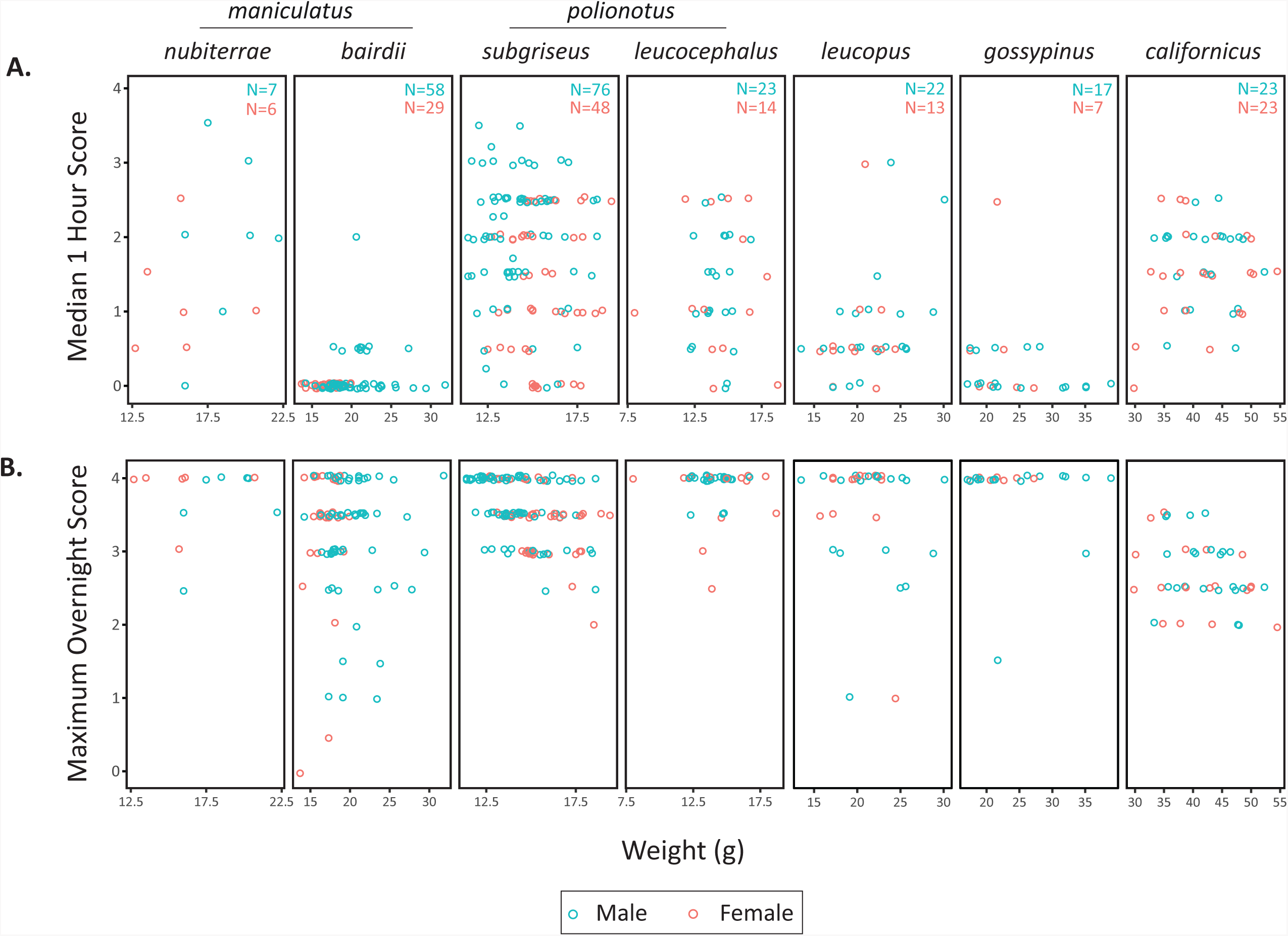
Effect of body weight on nesting behaviour by species and sex. (A) Median nest scores at 1h were not correlated with weight within any species-sex group (Spearman correlation, Bonferroni-corrected *P*>0.05). (B) Overnight maximum nest scores were not correlated with weight within any species-sex group (Spearman correlation, Bonferroni-corrected *P*>0.05). Note different x-axis for each species as indicated. Sample sizes are provided by sex and were the same for both time points.

**Supplemental Table S1:** Detailed Nest Scoring Criteria

| **Score** | **Score Description** | **Shredding** | **Nest Site<sup>1</sup>** | **Walls<sup>2</sup>** | **Overhead Cover** |
| --- | --- | --- | --- | --- | --- |
| **0** | **No Manipulation** | None | - | - | - |
| **0.5** | **Minor Shredding** | Minor:<br>≤top of 1 nestlet | - | - | - |
| **1** | **Extensive Shredding** | Extensive:<br>> top of 1 nestlet | No:<br>neither a nor b is true | No | - |
| **1.5** | **Ambiguous Nest Site** | Extensive | Unclear:<br>either a OR b is true | No | - |
| **2** | **Platform Nest** | - | Yes: both a and b are true | No: < ½ sphere height for < ½ circumference | - |
| **2.5** | **Partial Cup Nest** | - | Yes | Partial: < ½ sphere height for > ½ circumference OR ≥½ sphere height for < ½ of circumference | - |
| **3** | **Cup Nest** | - | Yes | Yes:<br>≥½ sphere height for ≥ ½ circumference | no overhead cover |
| **3.5** | **Partial Dome Nest** | - | Yes | Yes | Partial: <50% of the sphere is covered or there are multiple entrance holes |
| **4** | **Full Dome Nest** | - | Yes | Yes | Yes: ≥50% of the sphere is covered and there is at most one entrance hole |

1. A nest site is defined according to two criteria: **(a)** there is a contiguous concentration of nestlet around a central point consisting of ≥90% of any material the animal has shredded, and **(b)** the shape of the putative nest site is defined by the shredded material (>50% of the nestlet at the nest site is shredded). 2. To evaluate walls, imagine that the nest cavity is filled by a sphere, *sensu* (Hess et al., 2008). The walls are compared to the height of the sphere within the cavity, and the proportion of the circumference of the sphere that is surrounded by walls is noted.

**Supplemental Table S2:** Pairwise species comparisons of 1hr median and overnight maximum scores

| Species | *maniculatus nubiterrae* | *maniculatus bairdii* | *polionotus subgriseus* | *polionotus leucoceph.* | *leucopus* | *gossypinus* | *californicus (5g)* | *californicus (20g)* |
| --- | --- | --- | --- | --- | --- | --- | --- | --- |
| N: | 47 | 95 | 130 | 37 | 35 | 27 | 48 | 23 |
| *man. nub.* | | $W=3224.5$,<br>$P=9.3 \times 10^{-5}$ | $W=4124.5$,<br>$P=0.0026$ | $W=819.5$,<br>$P=1$ | $W=905$,<br>$P=1$ | $W=540$,<br>$P=1$ | $W=2102$,<br>$P=8.9 \times 10^{-13}$ | $W=974.5$,<br>$P=7.6 \times 10^{-8}$ |
| *man. baird.* | $W=4046$,<br>$P<2.2 \times 10^{-16}$ | | $W=5306.5$,<br>$P=1$ | $W=846$,<br>$P=1.8 \times 10^{-05}$ | $W=1133$,<br>$P=0.073$ | $W=545$,<br>$P=2.8 \times 10^{-5}$ | $W=3473$,<br>$P=3.8 \times 10^{-6}$ | $W=1553.5$,<br>$P=0.027$ |
| *pol. sub.* | $W=2565$,<br>$P=1$ | $W=734$,<br>$P<2.2 \times 10^{-16}$ | | $W=1353.5$,<br>$P=0.00025$ | $W=1778$,<br>$P=0.72$ | $W=866$,<br>$P=0.0017$ | $W=5435.5$,<br>$P=8.6 \times 10^{-14}$ | $W=2428.5$,<br>$P=1.3 \times 10^{-5}$ |
| *pol. leuc.* | $W=924.5$,<br>$P=1$ | $W=256.6$,<br>$P<2.2 \times 10^{-16}$ | $W=3088.5$,<br>$P=0.16$ | | $W=746.5$,<br>$P=1$ | $W=447.5$,<br>$P=1$ | $W=1693$,<br>$P=2.9 \times 10^{-12}$ | $W=791$,<br>$P=3.9 \times 10^{-8}$ |
| *leucop.* | $W=1159.5$,<br>$P=0.027$ | $W=375$,<br>$P=1.7 \times 10^{-15}$ | $W=3589.5$,<br>$P=2.5 \times 10^{-6}$ | $W=929.5$,<br>$P=0.023$ | | $W=362$,<br>$P=0.59$ | $W=1438.5$,<br>$P=2.9 \times 10^{-7}$ | $W=666$,<br>$P=0.00024$ |
| *gossy.* | $W=1095$,<br>$P=2.3 \times 10^{-6}$ | $W=1036$,<br>$P=0.29$ | $W=3176$,<br>$P=4.6 \times 10^{-10}$ | $W=881.5$,<br>$P=1.8 \times 10^{-6}$ | $W=763.5$,<br>$P=0.0002$ | | $W=1221$,<br>$P=1.2 \times 10^{-9}$ | $W=576$,<br>$P=4.9 \times 10^{-7}$ |
| *califor.* | $W=990.5$,<br>$P=1$ | $W=103$,<br>$P<2.2 \times 10^{-16}$ | $W=3492$,<br>$P=1$ | $W=678.5$,<br>$P=1$ | $W=281$,<br>$P=3.1 \times 10^{-6}$ | $W=74.5$,<br>$P=2.4 \times 10^{-9}$ | | |

The results of pairwise Wilcoxon rank-sum tests for species differences in median 1hr scores (below diagonal) or maximum overnight scores (above diagonal). For each comparison, test statistics (W) and Bonferroni-corrected p-values are reported; significant results ( $p<0.05$ ) are in bold.

**Supplemental Table S3:** Sex differences in nest scores

| **Taxon** | **Males** | **Females** | **1hr Median Score** | **Maximum Overnight Score** |
| --- | --- | --- | --- | --- |
| *maniculatus nubiterrae* | 31 | 16 | **W=119.5, P=0.03** | W=266, P=1 |
| *maniculatus bairdii* | 62 | 33 | W=858, P=0.11 | W=1173, P=1 |
| *polionotus subgriseus* | 80 | 50 | **W=1262.5, P=0.002** | **W=1361, P=0.008** |
| *polionotus leucocephalus* | 23 | 14 | W=163.5, P=1 | W=133, P=1 |
| *leucopus* | 22 | 13 | W=125.5, P=1 | W=152, P=1 |
| *gossypinus* | 19 | 8 | W=87, P=1 | W=84, P=1 |
| *californicus* | 25 | 23 | W=219.5, P=1 | W=238.5, P=1 |

The results of pairwise Wilcoxon rank-sum tests for sex differences in 1hr median nest scores or maximum overnight nest scores. Sample sizes, test statistics (W), and Bonferroni-corrected p-values are reported; significant results ( $P<0.05$ ) are in bold.

**Supplemental Table S4:** Spearman correlations between weight and nest scores within species/sex groups

| Taxon | Weight vs. 1hr Median Score |  | Weight vs. Maximum Overnight Score |  |
| --- | --- | --- | --- | --- |
|  | Males | Females | Males | Females |
| *maniculatus nubiterrae* | $r_s=0.24, N=7, P=1$ | $r_s=-0.09, N=6, P=1$ | $r_s=0.39, N=7, P=1$ | $r_s=0.13, N=6, P=1$ |
| *maniculatus bairdii* | $r_s=0.22, N=58, P=1$ | $N=29$, see below | $r_s=0.07, N=58, P=1$ | $r_s=0.13, N=29, P=1$ |
| *polionotus subgriseus* | $r_s=0.01, N=76, P=1$ | $r_s=0.04, N=48, P=1$ | $r_s=-0.32, N=76, P=0.08$ | $r_s=-0.31, N=48, P=0.48$ |
| *polionotus leucocephalus* | $r_s=0.03, N=23, P=1$ | $r_s=-0.10, N=14, P=1$ | $r_s=0.11, N=23, P=1$ | $r_s=-0.06, N=14, P=1$ |
| *leucopus* | $r_s=0.41, N=22, P=0.82$ | $r_s=0.22, N=13, P=1$ | $r_s=-0.08, N=22, P=1$ | $r_s=-0.04, N=13, P=1$ |
| *gossypinus* | $r_s=-0.25, N=17, P=1$ | $r_s=-0.24, N=7, P=1$ | $r_s=-0.23, N=17, P=1$ | $N=7$, see below |
| *californicus* | $r_s=-0.08, N=23, P=1$ | $r_s=0.14, N=23, P=1$ | $r_s=-0.36, N=23, P=1$ | $r_s=-0.29, N=23, P=1$ |

Sample sizes, Spearman correlation coefficient ( $r_s$ ), and Bonferroni-corrected p-values are reported. Sample sizes are smaller than for other tests due to missing weight data. We were unable to perform correlations between 1hr scores and weight within *maniculatus bairdii* females because all 29 animals received a median score of 0 at 1 hour. Similarly, all female *gossypinus* animals produced maximum scores of 4 at the overnight time point.

